# LAM – an image analysis method for regionally defined organ-wide cellular phenotyping of the *Drosophila* midgut

**DOI:** 10.1101/2021.01.20.427422

**Authors:** Arto Viitanen, Josef Gullmets, Jack Morikka, Pekka Katajisto, Jaakko Mattila, Ville Hietakangas

**Affiliations:** Faculty of Biological and Environmental Sciences, University of Helsinki, Finland; Institute of Biotechnology, University of Helsinki, Finland

**Keywords:** *Drosophila*, midgut, organ wide, imaging, intestinal stem cells, regeneration

## Abstract

Intestine is divided into functionally distinct regions along the anteroposterior (A/P) axis. How the regional identity influences the function of intestinal stem cells (ISCs) and their offspring remain largely unresolved. We introduce an imaging-based method, ‘Linear Analysis of Midgut’ (LAM), which allows quantitative regionally defined cellular phenotyping of the whole *Drosophila* midgut. LAM transforms image-derived cellular data from three-dimensional midguts into a linearized representation, binning it into segments along the A/P axis. Through automated multi-variate determination of regional borders, LAM allows mapping and comparing cellular features and frequencies with subregional resolution. Through the use of LAM, we quantify the distributions of ISCs, enteroblasts and enteroendocrine cells in a steady state midgut, and reveal unprecedented regional heterogeneity in the ISC response to a *Drosophila* model of colitis. Taken together, LAM is a powerful tool for organ-wide quantitative analysis of the regional heterogeneity of midgut cells.

**Graphical abstract:** 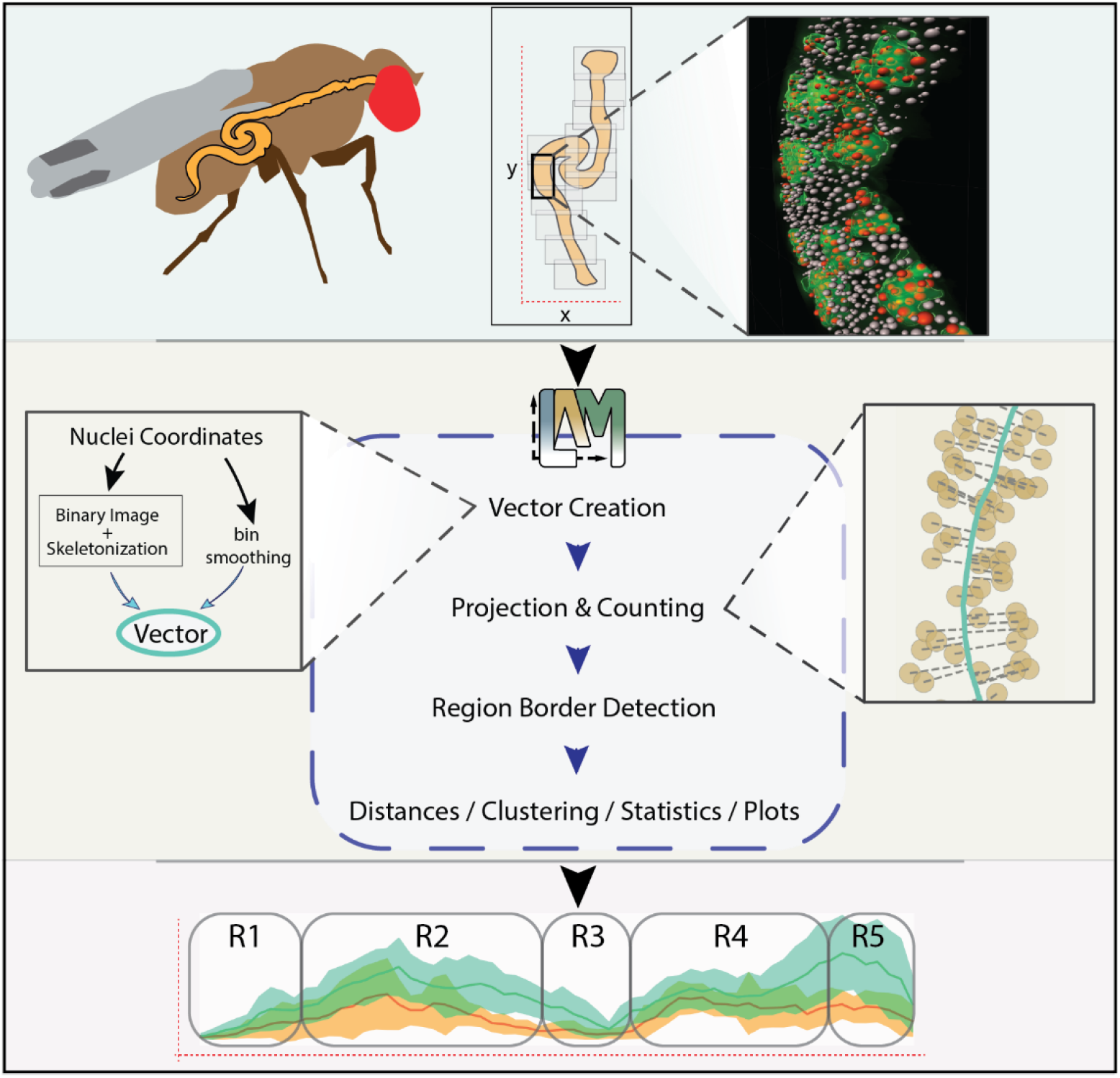

## INTRODUCTION

The intestine has a critical role in regulating organismal metabolism and immunity (Miguel-Aliaga et al., 2018). These functions are dynamically modulated by environmental factors, such as nutrition and microbes. Uncovering the mechanistic basis of the underlying regulation requires tractable *in vivo* model systems. The *Drosophila* midgut, analogous to the mammalian small intestine, has proven to be a powerful model for understanding intestinal physiology (Miguel-Aliaga et al., 2018). The midgut is composed of four cell types: the absorptive enterocytes (ECs), their differentiating progenitor cells, enteroblasts (EBs), the hormone secreting enteroendocrine cells (EEs), and the mitotic intestinal stem cells (ISCs) (Miguel-Aliaga et al., 2018). The midgut is an adaptive regenerative organ, whose cellular turnover and composition is affected by diet, sex, inflammation, age and reproductive status (Biteau et al., 2008; Buchon et al., 2009, 2013; Hudry et al., 2016; Reiff et al., 2015). Previous studies have uncovered regulatory pathways involved in the control of intestinal homeostasis through inter- and intracellular signaling (Gervais and Bardin, 2017; Guo et al., 2016).

In order to perform the functions of digestion, absorption, metabolism, nutrient sensing and signaling in a sequentially coordinated manner, the animal intestine is compartmentalized into regions along its anteroposterior (A/P) axis (Miguel-Aliaga et al., 2018; O’Brien, 2013). Moreover, human intestinal pathophysiologies, such as cancer or inflammatory disorders, often manifest in a region-specific manner (Missiaglia et al., 2014; Mowat and Agace, 2014). Therefore, the mechanisms that establish, maintain and modulate the regionalized functions of the intestine are of high biological and medical relevance. The *Drosophila* midgut regions have been distinguished based on anatomical characteristics, differential staining with histological dyes, as well as region-specific gene expression patterns (Buchon et al., 2013; Dimitriadis, 1991; Marianes and Spradling, 2013). Buchon et al. (2013) have divided the midgut into 6 major regions (R0-R5), which can be distinguished based on cross intestinal anatomy. R1-R5 were further divided into 14 subregions based on morphological, histological, and gene expression differences. In a parallel study, Marianes and Spradling (2013) divided the midgut into 10 zones, with significant overlap to the 14 subregions defined by Buchon et al. (2013).

Molecular analyses of the intestinal cell types have given more detailed insight into midgut regionalization. Consistent with sequentially coordinated digestion and absorption, the digestive enzyme and nutrient transporter genes display strictly region-specific expression patterns in the ECs (Dutta et al., 2015). The EE cells, mediating the signaling function of the intestine, can be divided into 10 subtypes displaying region-specific distribution (Guo et al., 2019). In addition to the differentiated cell types, it has also been proposed that the function of undifferentiated ISCs depends on regional identity. The ISCs display regional autonomy, i.e. their differentiated daughter cells do not cross most region boundaries (Marianes and Spradling, 2013). The ISCs in different midgut regions also display distinct morphological features, as well as differential gene expression, exemplified by the finding that more than 900 genes show regional expression variation in the ISCs (Dutta et al., 2015; Marianes and Spradling, 2013). The acidic R3 region, often termed the stomach of *Drosophila*, contains stem cells that have been deemed quiescent in unchallenged conditions, but activated in response to stressful stimuli, such as heat shock or pathogen ingestion (Strand and Micchelli, 2011). Despite the evidence strongly implying regional ISC heterogeneity, most studies on ISCs focus on one specific region (mostly R4) and the possible impact of regional identity and tissue environment on ISC regulation is largely overlooked.

Achieving representative data of the midgut requires unbiased quantitative analysis of all midgut regions. Rapid development of affordable and fast tile scan imaging has made it feasible to collect high resolution image data from the whole midgut. High phenotypic variation between midguts limits the reproducibility of qualitative analysis, and sets the requirement for robust quantitative analysis of replicate samples. However, achieving quantitative and regionally defined data from midgut cells has remained a major bottleneck, hampering the use of organ-wide analysis. Here we describe a widely applicable phenotyping method called LAM (Linear Analysis of Midgut) to achieve spatially defined quantitative data on midgut cells. As a proof of concept, we use LAM to quantitatively analyze regional distributions of ISCs, EBs and EE cells. We also demonstrate the regional heterogeneity of the regenerative response to a well-established colitis model, dextran sulfate sodium (DSS) treatment. The organ-wide analysis by using LAM revealed several novel features of DSS induced regeneration, including a failure of regenerative stem cell activation in R3, a regionally discordant pattern of stem cell division and differentiation in R4 vs. R5, as well as an increase in enteroendocrine cell numbers in the posterior R4/R5 region. By making unbiased, quantitative, organ-wide analysis highly feasible, LAM is expected to open new avenues for the analysis of regional heterogeneity of midgut cells.

## DESIGN

Because of its small size and compartmentalized structure, the midgut is an ideal model for performing tissue-wide analyses to address the role of regional identity and tissue environment for the function and regulation of individual intestinal cells. Although imaging of whole midguts is feasible by using affordable and fast tile scan imaging, several biological features of the midgut hamper the downstream analysis. The midguts have a coiled structure and their intraluminal content is variable. Therefore, each midgut has a unique morphology, which brings about the need for time-consuming and subjective manual work in order to identify, align and compare the intestinal regions. In order to perform these functions in an automated and unbiased way, we developed means to transform the three-dimensional midgut images into one dimension. This was achieved by designing an algorithm enabling midline vector creation along the A/P axis. Coupling of cellular identities into a specific position on the linear vector enables binning of cell-specific data along the midgut. The one-dimensional data enabled automatization of the identification of regional boundaries, allowing accurate aligning of corresponding regions. These features enabled LAM to achieve robust quantitative phenotyping of midguts with subregional resolution. The quantification approach of LAM produces large and information-dense datasets. To facilitate the downstream data analysis, we have included various options for visualization, statistical analysis, and data subsetting. A graphical user interface, user manual and tutorial videos make LAM accessible to all researchers.

## RESULTS

### An approach for spatially defined quantitative phenotyping of the Drosophila midgut

In order to analyze the spatial heterogeneity of intestinal cell responses in an unbiased and reproducible manner, we developed an intestinal phenotyping approach, which is automated, quantitative, and regionally defined. For imaging the nuclei of pseudostratified midgut epithelium, fixed DAPI stained tissues were mounted in between a coverslip, and a microscope slide with 0.12 mm spacers. Flattening the intestinal tube into two epithelial layers, yet separated by its lumen, allowed Z stack acquisition of one layer, saving time and reducing file size (Fig 1A). As an initial step, we sought a means to reduce the tile scan stacks of non-linear midguts into a linearized representation (Fig 1B). For this, an algorithm that determines the midline vector along the A/P axis was used. Nuclei, whose positions were plotted along X:Y:Z co-ordinates, were projected onto this midline vector (Fig 1C). Consequently, the positional information from the X:Y:Z co-ordinates could be reduced to one dimension, i.e. to a position on the midgut A/P axis. The vector was then divided into bins, the number of which can be adjusted to a desired spatial resolution. Due to the positional information and binning of the measured cellular features, data collected from different intestines could be aligned and combined within a data matrix as biological replicates, and compared between sample groups, thus allowing quantitative representation and statistical analysis.

**Figure 1.**
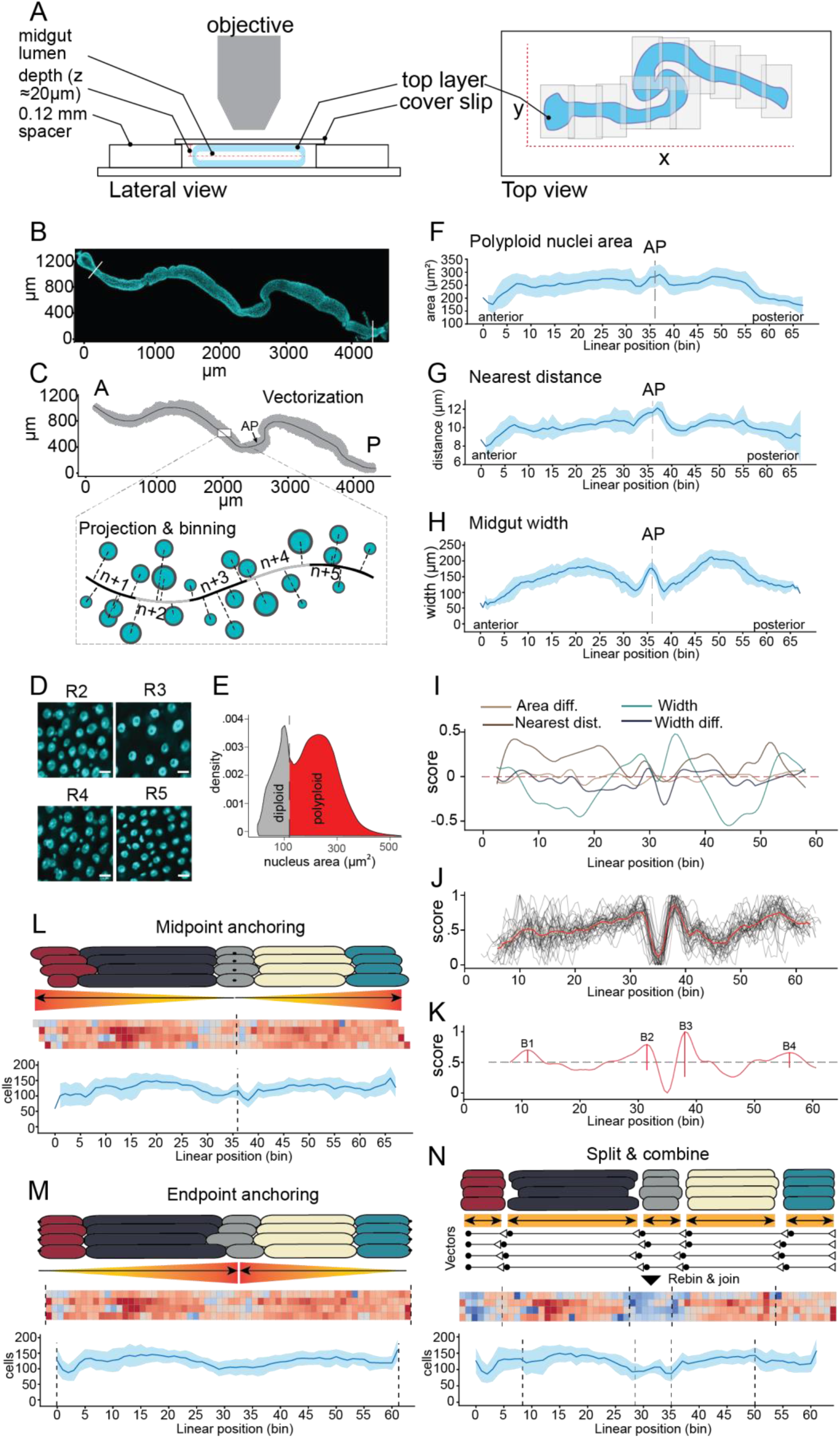
A pipeline for regionally defined quantification of *Drosophila* midguts. **A)** Schematic presentation of the whole midgut imaging. **B)** A representative tile scan image of DAPI (cyan) stained midgut. After imaging and stitching of the tiles, the image is processed to exclude any features lying outside the area of interest. Subsequently, the image is analyzed for DAPI spots by, for example, the spot detection algorithm of the Imaris software. **C)** Vector building, projection and linearization. The X:Y co-ordinates of the nuclei centroids are utilized to build a vector along the midline of the midgut image. The spots, and any accompanying data, are then projected onto the vector. The vector is then binned, where the number of bins is chosen based on the desired resolution. An “anchoring point” (AP) is introduced into a morphologically distinct place, such as the border between the copper cell region (CCR) and the large flat cells (LFC) region of the middle midgut (arrow). **D)** Midgut regions have distinct enterocyte size and density. Representative images of DAPI (cyan) stained nuclei from the midgut R2-R5 regions. Scale bar 10 µm. **E)** Nuclei area profile from whole midguts determined by identification of DAPI spot areas from the Imaris spot detection algorithm. For the subsequent analysis of midgut region borders, the diploid cells were filtered out from the dataset. **F)** Polyploid nuclei area profile along the A/P axis of midgut. AP=Anchoring Point. N=32 midguts. Light blue shading is the standard deviation. **G)** Nuclei nearest distance profile along the A/P axis of midgut. The distance between nuclei is a proxy for cell density. AP=Anchoring Point. N=32 midguts. Light blue shading is the standard deviation. **H)** Width profile along the A/P axis of midgut. Midgut width is approximated by following the vector bin-by-bin and summing the average projection distances of the most distant decile of cells on both sides of the vector. AP=Anchoring Point. N=32 midguts. Light blue shading is the standard deviation. **I-K)** Border detection algorithm performs a multivariate border region detection for each sample and outputs average border locations for each sample group. **I)** Smoothed scores of default border detection variables along A/P axis of a sample. The variables are scored based on weighted divergence from expected values, i.e., from a fitted fifth degree Chebyshev polynomial. The variable scores are summed to provide a total score for each location of a sample, which are then re-scaled to interval [0, 1] in order to give comparable peak locations despite phenotypic differences. **J)** Total scores of samples belonging to one sample group (N=32). The red line is the median score of the sample group and black lines are individual samples. While individual samples have great variation in score, grouping of samples leads to emergence of trends that can be used for peak detection. **K)** Peak detection performed on a sample group’s median scores (red line) shows approximate locations of border regions, as defined by value changes in multiple variables. The group’s score is smoothed and re-scaled to [0, 1] for peak detection. The vertical red lines at peak locations show their prominences. The marked borders from left to right are B1, B2, B3 and B4. **L-N)** Anchoring of midgut samples for regional alignment. **L)** Midpoint anchoring. Using a single anchoring point in a distinct morphological site, such as the border between Cu cells and large flat cells, results in accurate alignment close to the anchoring point but propagates error toward the distal regions. The anchoring point is a user-defined image co-ordinate that is projected onto the normalized [0,1] vector. The vector is then divided into a user defined number of bins that is equal for each sample. The samples are aligned within a data matrix by assigning them to indices according to the bin of their projected anchoring point. Note the unequal alignment of the midgut ends due to varying proportions of regions, and variable lengths at either side of anchoring point. **M)** Endpoint anchoring. Aligning the samples from both ends propagates the error towards the middle of the midgut. In this method, a user defined anchoring point is not necessary. **N)** Split and combine anchoring. In this method, border peak analysis determines vector cut points. This allows splitting, realigning, and rejoining of the vectors with accurate regional comparison of different midgut samples.

Next, we wanted to address, whether the quantitative information obtained by using our algorithm allowed automatic determination of the borders of midgut regions. Midgut regions are characterized by differences in enterocyte ploidy, and density (Fig 1D) (Marianes and Spradling, 2013) and they are separated by constrictions of the midgut radius (Buchon et al., 2013). We first separated the polyploid enteroblast/enterocyte population from diploid cells based on nuclear area (Fig 1E). The data on polyploid nuclear area, nucleus-to-nucleus distance as well as midgut width was then projected onto the A/P vector. Due to a high variability of morphology, it was not possible to reliably detect region borders from individual midguts (data not shown). This led us to explore the border detection from combined measurements of several replicate samples. In order to align the data of individual midguts for bin-to-bin comparisons, we introduced an anchoring point (AP) in the middle of the midgut, located at the border of the copper cell (CCR) and large flat cell (LFC) regions, which can be easily identified based on the difference in nuclear distance (Fig 1C). Combining the measurements of polyploid nuclear area, nucleus-to-nucleus distance as well as midgut width from several replicate midguts revealed characteristic midgut profiles, as described by Buchon et al., 2013 (Fig 1F-H). A multivariate border detection algorithm based on these profiles allowed robust identification of four peaks corresponding to region borders, B1-B4 (Fig 1I-K).

Midgut total length, as well as the length of the individual regions, is variable (Buchon et al., 2013). This poses a challenge for aligning corresponding regions of replicate samples. Accordingly, the utilization of a single alignment point in the middle midgut, i.e. the point where the vectors of different samples are anchored together, can lead to an imprecise alignment of the regions towards the anterior and posterior ends (Fig 1L). On the other hand, anchoring the samples from the ends will reduce the accuracy of the alignment in the middle regions (Fig 1M). To minimize noise introduced by the variable length, we utilized region border analysis to apply several independent alignment points resulting in an optimal comparison of midguts regions. In this “split and combine” approach the vectors were cut based on region border detection, aligned separately, and rejoined back together (Fig 1N). This pipeline improved accuracy in regional comparisons between the midguts.

We have implemented the above-described analysis tools into a python package, called “Linear Analysis of Midgut” or LAM (https://github.com/hietakangas-laboratory/LAM). LAM provides various options for analyzing midgut image derived feature data, such as object co-ordinates for measuring cell-to-cell distances and cell clustering, object size, and object intensities in a regional manner. It also provides various options for plotting and statistical analysis between sample groups. We also provide a separate tool for stitching tile images for large scale data sets (https://github.com/hietakangas-laboratory/Stitch). Finally, LAM is accompanied by a step-by-step guide, tutorial videos, as well as a user-friendly graphical user interface.

### Region-specific cellular profiling of the steady-state Drosophila midgut

To date, no quantitative data on regional distribution of cell types in a steady-state midgut is available. We used LAM to establish such a dataset for mated young (7d) females, grown on chemically-defined holidic media (Fig 2A) (Piper et al., 2014). With the chosen experimental settings, we expect the midguts to be in a gradually renewing steady state. The border detection algorithm was used to identify regions R1-R5. To identify intestinal cell types, specific markers for ISCs (Delta-LacZ), EBs (Su(H)-LacZ), and EEs (anti-Prospero) were used along with Esg-Gal4,UAS-GFP,tub-Gal80^ts^ (Esg^ts^), which marks ISCs and EBs (Fig S1 A, B) (Jiang et al., 2009). The total and relative (normalized to total cell number) cell numbers within regions R1-R5 were calculated (Fig 2B-D and Fig 1S C-F). The analysis shows clear regional variation in the proportional numbers of distinct cell types, for example, the EE cells were most concentrated in R3 (Fig 2D). The overall regional pattern of Delta positive ISC and Su(H) positive EB distributions largely overlap with each other (Fig 2E, F). The relative number of ISCs and EBs were high in the middle, and the posterior of R4 (corresponding to R4bc) as well as in the anterior R5 (corresponding to the R5a). In R2, ISCs and EBs are most abundant in the middle of the region (corresponding to the R2b). Notably, R1 contains very low numbers of ISCs and EBs, compared to the rest of the midgut (Fig 2E, F).

**Figure 2.**
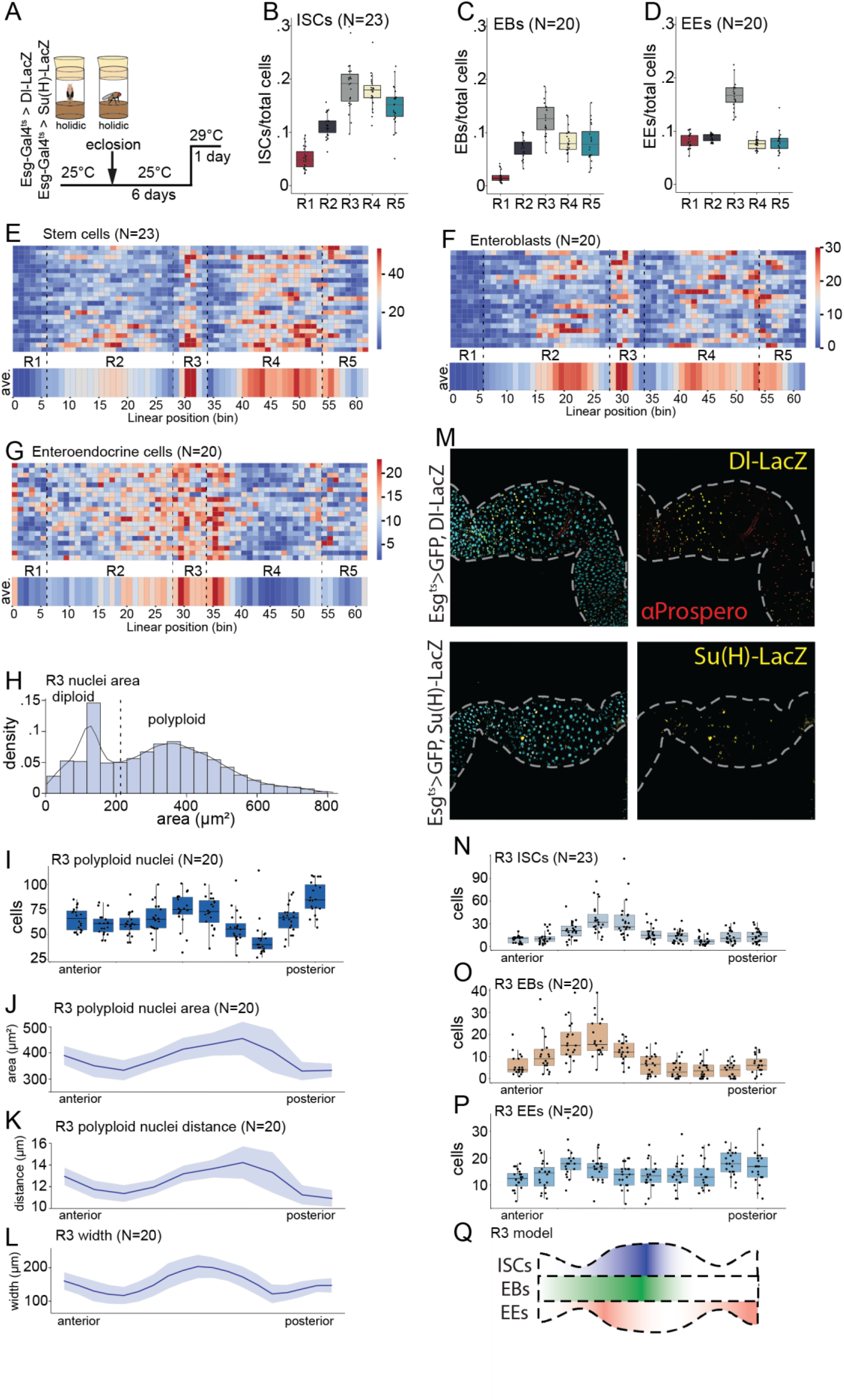
Cellular profiling of a steady state midgut by LAM. **A)** Experimental design used for the regional steady state midgut profiling. Age matched, mated females of Esg-Gal4^ts^, UAS-GFP > Delta-LacZ or Esg-Gal4^ts^, UAS-GFP > Su(H)-LacZ genotype were kept at +25°C for six days, and then shifted to +29°C for one day. **B-D)** Relative numbers of ISCs (B) EBs (C) and EEs (D) in R1-R5 regions. Calculated as number of specific cells per total number of cells. **E-G)** Sample and average heat maps of cellular distributions along the midgut A/P axis for ISCs (E) EBs (F) and EEs (G). **H)** Area distribution of R3 nuclei. **I)** Polyploid nuclei number along the R3 A/P axis. **J)** Polyploid nuclei area along the R3 A/P axis. Light blue shading is the standard deviation. **K)** Average polyploid nuclei to nuclei distance along the R3 A/P axis. Light blue shading is the standard deviation. **L)** R3 width along the A/P axis. Light blue shading is the standard deviation. **M)** Representative images of R3 region showing the localization of Dl-lacZ-positive ISCs and Prospero-positive EEs (upper panels) and Su(H)-lacZ-positive EBs (lower panels). DNA is stained with DAPI and is shown in cyan. **N)** ISC number along the R3 A/P axis. **O)** EB number along the R3 A/P axis. **P)** EE number along the R3 A/P axis. **Q)** A schematic model deciphering the steady state distribution of ISCs, EBs and EEs in the R3 region.

As the LAM analysis was performed at the resolution of 62 bins/midgut, we were able to identify even more fine-structured patterns of cellular distribution. For example, R3 is divided into the acid secreting copper cell (CCR), and the large flat cell (LFC) regions flanked by intestinal constrictions. Plotting the polyploid EC nuclei number, area, nuclei to nuclei distance, and midgut width revealed typical topology of the CCR and LFC along the R3 region (Fig 2H-L) Interestingly, the anterior side of R3, composed of the CCR, displayed high relative numbers of ISCs and EBs. However, their respective distributions within this region differed slightly: ISCs were most abundant in the middle/posterior part of CCR, while EBs were primarily clustered in the anterior end of the CCR, adjacent to the R2/R3 border (Fig 2M-O). This is in line with the findings that the CCR can be subdivided into molecularly distinct regions (Strand and Micchelli, 2011) and suggests the existence of localized signals directing the balance between stem cell renewal and differentiation in the CCR. In addition to ISCs and EBs, EE cells displayed specific patterns in the middle midgut (Fig 2M,P & S). A high density of EEs was present in a narrow stripe at the anterior CCR, as well as right after the R3/R4 border (Fig 2P). The latter stripe corresponded to the so-called iron cell region, which contains enterocytes highly expressing the iron storage protein Ferritin (Marianes and Spradling, 2013). Interestingly, additional enrichments of the EE cells were observed in the distal ends of the midgut, at the border between the crop and R1, and at the border between the midgut and hindgut (Fig 2G). Taken together, profiling of the cellular distributions along the steady-state midgut A/P axis by LAM revealed unprecedented patterns of cell organization, and demonstrated the performance of LAM in quantitative analysis of sub-regional phenotypic features (Fig 2Q).

### DSS feeding results in regional changes to midgut morphology and ISC differentiation

As a further proof-of-concept of the functionalities in LAM, we analyzed the regenerative response of ISCs in a widely used colitis model, oral administration of dextran sulfate sodium (DSS) (Fig 3A, B). DSS treatment has been reported to induce regenerative ISC proliferation and accumulation of Su(H) positive enteroblasts, with no significant changes in numbers of Delta or Prospero positive cells (Amcheslavsky et al., 2009). An analysis of the morphological features of the midgut revealed that DSS feeding results in significant, region-specific changes in midgut morphology. Midgut width, and length were affected in several regions, especially in R3 and R4 (Fig 3C-D). Furthermore, the size and patterning of nuclei were altered in a region-specific manner (Fig 3E). These changes somewhat compromised border detection, in particular preventing reliable detection of the first border (B1, Fig 3F). The most striking morphological consequence of the DSS feeding was the reduction of R3 size (Fig 3D, G). This suggests that the ISCs of R3 are not capable of maintaining homeostatic regeneration upon DSS treatment. Accordingly, in R3, the DSS treatment resulted in significant loss of enterocytes, whereas the number of smaller diploid nuclei was less affected (Fig 3H). In line with this result, the number of ISC-derived GFP marked cells were not significantly increased in the R3 region upon DSS treatment, consistent with the notion of the stem cells’ inability to divide and compensate for cell death (Fig 3I). As a consequence, the typical subregional R3 morphology, including differential patterning and number of ECs in the CCR and LFC regions, was lost in the DSS treated flies (3J, K). Taken together, based on the analysis of the morphological and cellular parameters, our results show the severe inability of stem cells in the R3 region to compensate the cell loss upon DSS treatment.

**Figure 3.**
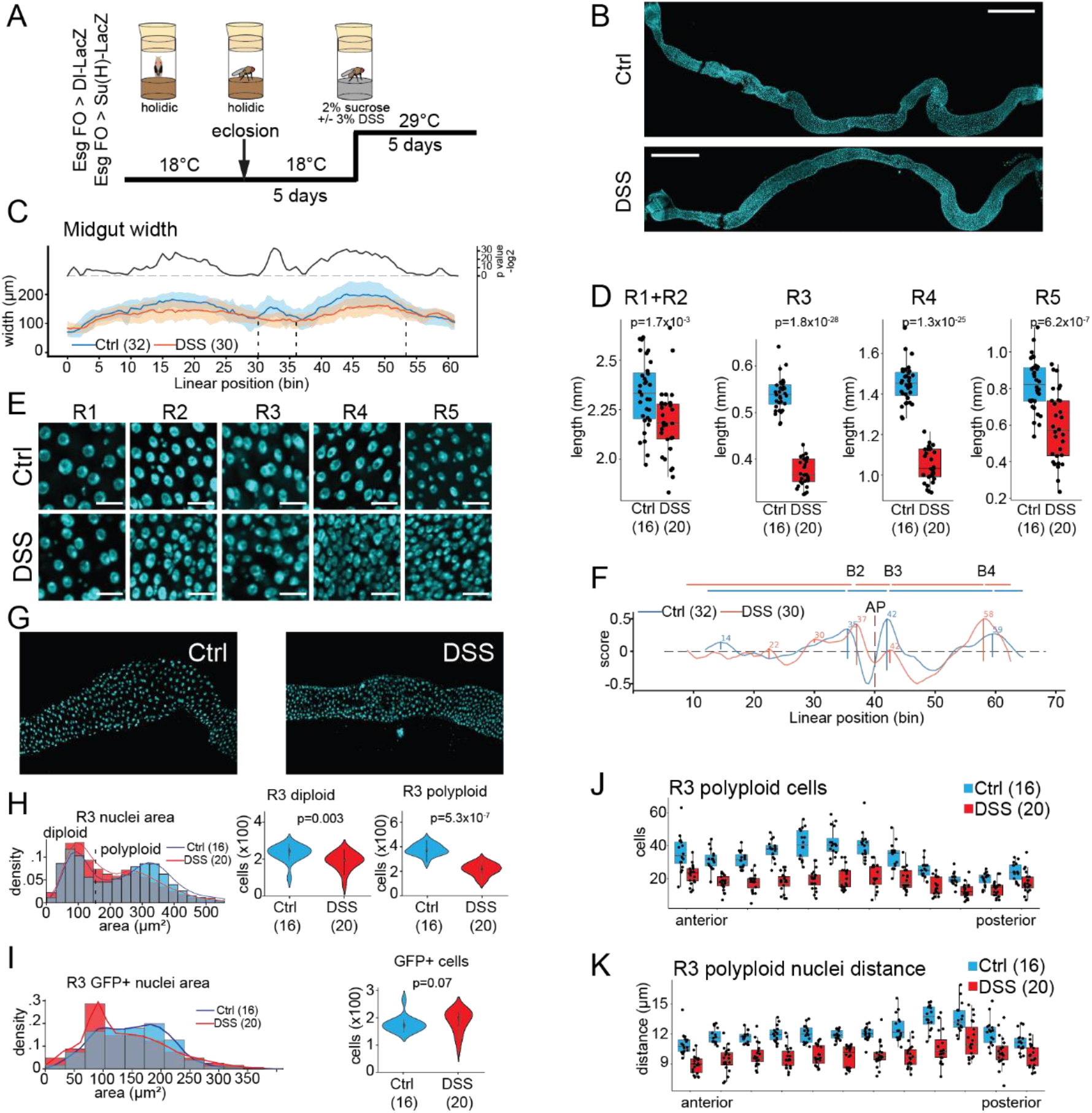
DSS feeding results in regional changes to midgut morphology. **A)** Experimental design of the DSS feeding experiment. Age matched, mated females of Esg FO > Delta-LacZ or Esg FO > Su(H)-LacZ genotypes were kept at the restrictive temperature (+18°C) for 5 days, and then shifted to the permissive temperature (+29°C) to induce the flip out clones in the presence of 3% DSS. **B)** Representative images of midguts of control and DSS fed flies. DNA is stained with DAPI and is shown in cyan. Scale bar 500 µm. **C)** Width profile of the control and DSS treated midguts. Light blue and orange shadings are the standard deviations. **D)** Length of R1+R2, R3, R4, and R5 midgut regions of control and DSS fed flies. **E)** Representative images of R1-R5 midgut regions of control and DSS fed flies. DNA is stained with DAPI and is shown in cyan. Scale bar 20 µm. **F)** Border peak analysis of midguts of control and DSS fed flies. **G)** Representative images of the R3 region of control (left panel) and DSS (right panel) fed flies. DNA is stained with DAPI and is shown in cyan. **H)** Area distribution of R3 nuclei in midguts of control and DSS fed flies. Comparison of the number of R3 diploid, and polyploid nuclei between midguts of control and DSS fed flies. **I)** Area distribution of R3 GFP positive nuclei in midguts of control and DSS fed flies. Comparison of the number of R3 GFP positive nuclei between midguts of control and DSS fed flies. **J)** Number of polyploid nuclei in R3. Anterior to the left, posterior to the right. **K)** Mean distance between polyploid nuclei in R3. Anterior to the left, posterior to the right. p values in D are calculated by the two-sample t-test. p values in H & I are calculated by Mann-Whitney-Wilcoxon U-test.

To further investigate the regional heterogeneity of ISC differentiation during DSS-induced regeneration, cell type specific markers for ISCs (Delta-LacZ), EBs (Su(H)-LacZ), and EEs (anti-Prospero) were used. Consistent with earlier findings (Amcheslavsky et al., 2009), the DSS treatment led to accumulation of Su(H) positive EBs (Fig 4A, B). However, the accumulation of EBs displayed region-specific differences, being most prominent in R5 and particularly low in R1, and in the anterior parts of R4 (Fig 4B). In contrast to the previous report (Amcheslavsky et al., 2009), we detected widespread accumulation of Delta positive ISCs, especially in R2 and R4 (Fig 4C, D). Notably, the regional pattern of Delta and Su(H) positive cells did not fully correlate, indicating that the symmetric ISC-ISC divisions (high Delta) are dominating in the R4, whereas asymmetric ISC-EB divisions (high Su(H)) are more prominent in R5 (Fig 4B, D). Interestingly, we also noticed that the nuclei of the Su(H) positive cells were larger in R5 compared to R4, suggesting a defective enteroblast to enterocyte differentiation in the R5 region (Fig 4A, E). Consistent with the low amount of stem cells in R1 during steady state, few Delta positive cells were detected in the anterior parts on the midgut following the DSS treatment (Fig 4D). The levels of Prospero positive EE cells remained stable upon the DSS treatment in most of the midgut area (Fig 4F). Interestingly, however, an area ranging from posterior R4 to anterior R5 displayed significantly elevated numbers of Prospero positive cell after the DSS treatment (Fig 4F). In conclusion, the DSS-induced regenerative response displays prominent regional heterogeneity, in terms of stem cell activation, division, as well as differentiation profiles.

**Figure 4.**
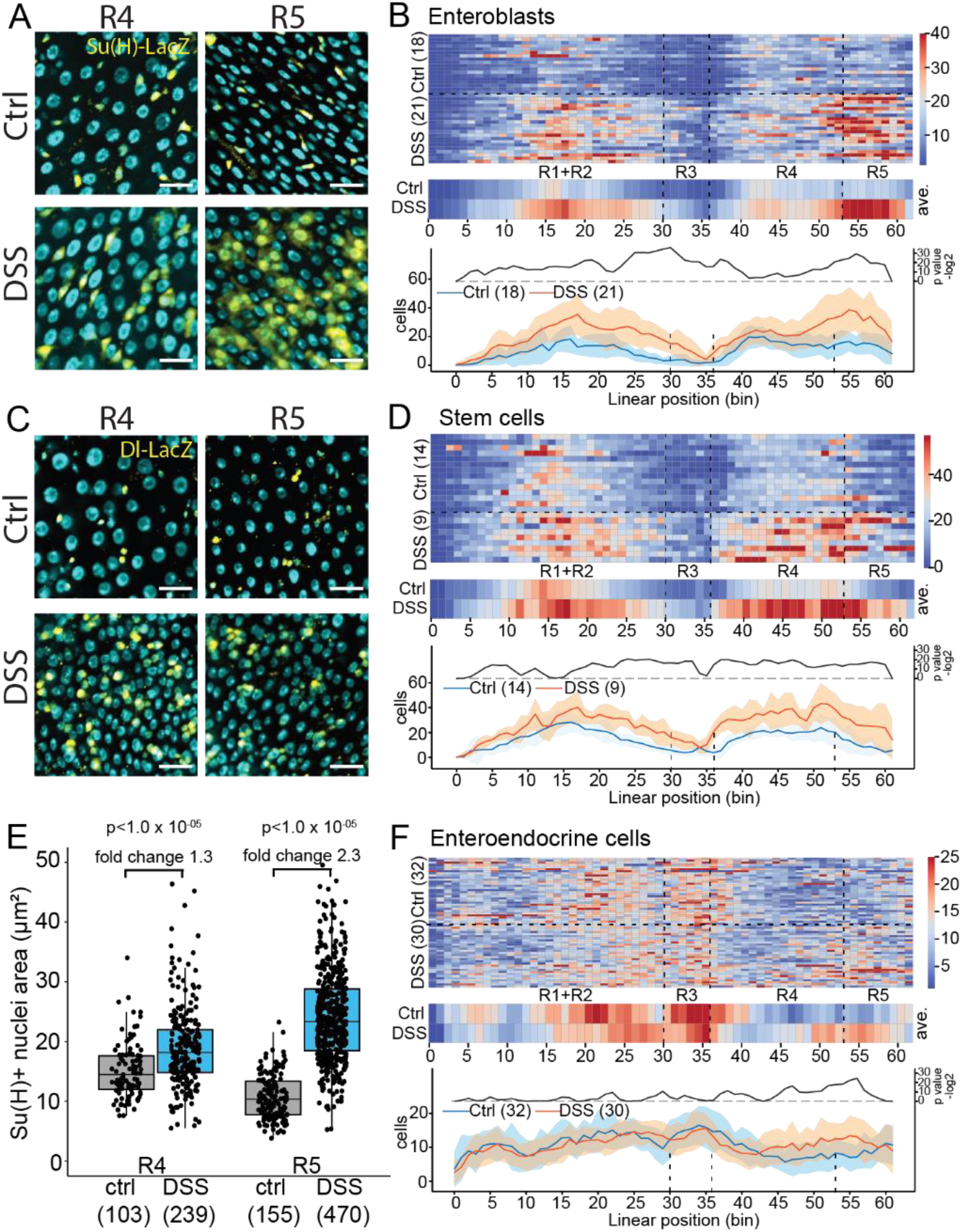
DSS feeding results in regional changes to ISC differentiation. **A)** Representative images of the R4 (left panels) and R5 (right panels) regions of ctrl and DSS fed midguts from Esg-Gal4^ts^ > Su(H)-LacZ flies. DNA is stained with DAPI and is shown in cyan. Scale bar 20 µm. **B)** Regional quantification of Su(H)-LacZ positive EBs of midguts of control and DSS fed flies. **C)** Representative images of the R4 (left panels) and R5 (right panels) regions of ctrl and DSS fed midguts from Esg-Gal4^ts^ > Delta-LacZ flies. DNA is stained with DAPI and is shown in cyan. Scale bar 20 µm. **D)** Regional quantification of Delta-LacZ positive ISCs of midguts of control and DSS fed flies. **E)** Nuclei area quantification of Su(H) positive cells in R4 and R5 regions. **F)** Regional quantification of Prospero positive EE cells of midguts of control and DSS fed flies. p values in E are calculated by two-way ANOVA followed by Tukey’s test.

## DISCUSSION

Here we present an approach to quantitatively study cellular phenotypes of the whole *Drosophila* midgut. In combination with fast tile scan imaging and efficient image feature detection algorithms, LAM enables, for the first time, quantitative and regionally defined automated phenotyping of all cells in the whole midgut. LAM allows (i) coupling of cellular identities to a specific position along the A/P axis, (ii) automated detection of regional boundaries, and consequently (iii) quantitative and statistical analysis of cellular phenotypes along the regions of the midgut, with sub-regional resolution. In doing so, LAM (iv) opens the path for organ-wide studies of midgut cells and eliminates the bias caused by a selective analysis of a specific midgut area. Through these advances, LAM will allow the exploration of regional heterogeneity of midgut cells, including the ISCs, and will significantly increase the representativeness of midgut phenotypic data. The graphical user interface makes LAM accessible even for scientists with limited experience in computational image analysis.

### Limitations of LAM, and potential solutions

The performance of LAM is dependent on the quality of the midgut preparations, image acquisition, and image segmentation for cellular objects. Each step is to be carefully considered for successful application of LAM. In our experiments, the intestines were mounted between a microscope slide with 0.12 mm spacers, and a coverslip. Images were obtained to capture half of the midgut circumference, thus assuming the cellular heterogeneity to be equal on each side. When mounted, the midgut is not always equally flat along the A/P axis. Special care is needed to avoid disproportional recording of the midgut circumference in different regions. To circumvent any bias from disproportional imaging, it is possible to extend the Z stacks to include the full circumference of the midgut if required. While LAM allows the recording of all imaged cells, the projection of objects to the midline vector is dimensionally restricted and LAM does not account for orientation on the Z-axis. Consequently, the information on cell stratification as well as the three-dimensional geometry of the intestinal cylinder is not used during object counting. Z-axis co-ordinates are, however, taken into account when calculating the object distances and clustering, allowing reliable data acquisition around the intestinal circumference.

The algorithm for detecting region borders is based on the morphological characteristics of the regions, such as midgut constrictions, as well as the nuclear size and distance, which were previously applied to manually map borders between physiologically distinct compartments (Buchon et al., 2013). Object segmentation is a critical step for calculating the nuclear features characteristic for different regions. The performance of the traditional object segmentation methods, such as intensity thresholding, is compromised by high cellular densities, cell size variation, cell stratification, and intensity differences. The variation in the success of nuclear segmentation possibly hampered our attempts to reliably detect regional borders from individual midguts. We overcame this limitation by applying sample group average values to locate the borders of individual samples. Deep learning based nuclear segmentation algorithms, such as StarDist (Schmidt et al., 2018; Weigert et al., 2020), are likely to further improve the accuracy.

### Novel features of regional distribution of midgut cells

We tested the performance of LAM by analyzing the distribution of ISCs, EBs and EE cells in a steady-state midgut of mated young females. This analysis revealed several new features of cellular distributions, including partially overlapping clusters of Delta and Su(H) positive cells within the CCR/R3ab subregion. In addition, EE cells were observed to cluster around the main regional boundaries, including the cardia-R1, R2-R3, R3-R4 and R5-hindgut boundaries, suggesting a common regional organizer for the specification of EE cell fate in these regions. One such signal could be the Wg signaling pathway, whose activity has been shown to localize to these regions (Tian et al., 2016). Notably, due to the high variation of phenotypes between individual midguts, it would have not been possible to reliably detect such features by qualitative analysis of individual midguts. This demonstrates the ability of LAM to detect variable phenotypes with high subregional resolution. The resolution of LAM is influenced by the numbers of bins, which can be freely adjusted by the user. The optimal number of bins depends on the density of input data points as well as data quality, which influences the accuracy of alignment of individual midguts.

### Regional heterogeneity of the regenerative response in a Drosophila colitis model

As another proof of principle, we employed a widely used *Drosophila* colitis model, induced by DSS feeding. Use of LAM allowed us to identify several new features of stem cell activation and differentiation not previously documented in the literature, providing new insight into the previously reported models of midgut regeneration (Amcheslavsky et al., 2009; Jiang et al., 2016). We noticed that the stem cells in R3 were markedly less capable of regenerating EC loss, when compared to the stem cells in the flanking R2 and R4 regions. This led to a significant reduction of the total cell numbers in R3. Thus, for the first time, our study shows that the gastric stem cells cannot efficiently regenerate lost tissue when challenged. DSS treatment led to a relatively uniform increase in Delta positive cells, except in R1, which contains fewer Delta-positive cells to start with. While the overall pattern of Su(H)-positive cells was similar to that of the Delta positive cells, the posterior midgut displayed interesting differences. Most of R4 showed only a modest increase in Su(H) positive cells, but an area from the posterior end of R4 to R5 displayed a very high increase in relative numbers of EBs in response to DSS. This suggests that the differentiation rate of ISCs displays significant regional differences, with the ISCs of R5 being more prone to EB fate. In addition, the Su(H) positive cells in the R5 region of the DSS treated midguts showed enlarged nuclei compared to the EB cells in other regions. While the molecular details explaining the difference in ISC fate between R4 and R5 are as yet unknown, regional transcriptome mapping has revealed existing gene expression differences between the ISCs of these regions (Dutta et al., 2015). One candidate in regulating ISC fate in these regions is the transcription factor Snail, whose expression is relatively high in R5 ISCs. Forced expression of Snail prevented EB differentiation into ECs, leading to an accumulation of EBs (Dutta et al., 2015). Hence, it will be interesting to learn whether intrinsic differences in Snail expression, or possible region-specific extrinsic factors, underlie the region-specific differentiation patterns in the regenerating midgut.

### Concluding remarks

In addition to the physiology of midgut regionality, the unbiased organ-wide analysis with LAM can improve representativeness of midgut data in general. Considering the real concern of confirmation bias throughout the scientific literature, there is a risk that studies focusing on a narrow (often undefined) area of the midgut primarily record and present data from areas that give the strongest phenotypes. Considering our DSS experiment, a focused analysis of only one (sub)region would have yielded several different, and sometimes even mutually contradictory, biological conclusions - depending on the region chosen. Therefore, one should exercise caution when making generalized conclusions based on the findings of a small subset of ISCs. We propose an approach where the phenotypic response for a given treatment/genotype is first quantitatively analyzed, and reported, at the level of the whole midgut, with more detailed follow-up experiments concentrating on the specific region(s) of interest. In conclusion, we expect that the unbiased organ-wide analysis offered by LAM will allow the pursuit of more representative data, and uncover the extent of tissue context-dependency of stem cell regulation as well as increasing understanding of the physiological roles of intestinal regionalization.

## MATERIALS & METHODS

### LAM

Data handling in LAM is performed with NumPy (Harris et al., 2020), and Pandas (McKinney, 2010), while plotting is done using matplotlib (Hunter, 2007) and Seaborn (Waskom et al., 2020). Geometric and image operations are performed with Shapely (Gillies et al., 2007) and Scikit-image (van der Walt et al., 2014), respectively. Statistics are calculated with scipy.stats (SciPy 1.0 Contributors et al., 2020) and statsmodels (Seabold and Perktold, 2010). The border detection additionally uses scipy.signal (SciPy 1.0 Contributors et al., 2020) for locating regions of high signal. LAM includes an easy-to-use graphical user interface (GUI) with enabling/disabling of related options as well as a default settings file that can be edited at will to control all runs. LAM also supports execution from command line using a limited scope of arguments. Full description of the usage of LAM, and step-by-step instructions can be found in the LAM user guide found in GitHub (https://github.com/hietakangas-laboratory/LAM). LAM video tutorials are available in (https://www.youtube.com/playlist?list=PLjv-8Gzxh3AynUtI3HaahU2oddMbDpgtx).

### Vector creation

LAM provides two alternative methods for creating piecewise median lines, which we colloquially call vectors, for midgut images: bin-smoothing and skeletonization. The methods require the midguts to be horizontally oriented. An auxiliary script is provided to rotate data to horizontal orientation. Bin-smoothing of the data is performed by binning the x-axis after which the median of the nuclei coordinate is calculated for each bin. Then a piecewise line is created to connect the bin midpoints. The number of bins is a user defined parameter to be adjusted for suitable level of smoothing. In the skeleton vector creation option the DAPI channel image is first converted into a binary image where each nuclei is resized to one pixel. The binary image is processed with resizing, smoothing, binary dilation, as well as hole filling in order to produce a continuous blob (user defined parameters, see user guide for more details). A binary matrix is then created where pixels of nuclei are marked as one, and empty pixels as zero. The matrix is then subjected to skeletonization, where pixels of the image are eroded until reduced to pixel-wide structures. The vector starting point is determined as the average of five pixel co-ordinates having the smallest x value. The vector is then drawn from pixel to pixel by scoring pixels within a specified range (find distance in GUI) using the following penalty function:

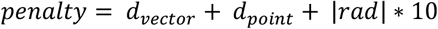

where *d_vector_* is the distance of the pixel to the last co-ordinate of the vector, and *d_point_* is the pixel’s distance to the projection point ahead of the last coordinate. The projection point is determined by adding the previous vector progression (distance and direction) into the last vector point. The final scoring component, the modulus of radians, is the difference in direction between the last vector co-ordinate and a pixel compared to a fitted line drawn between the last three vector co-ordinates. The x and y co-ordinates of the pixel with the smallest penalty are then added to the path of the vector, and the next pixels are scored based on these coordinates, and so on until no more pixels are found (Fig S2 A)

### Projection and counting

All segmented image objects, which we colloquially call features, and their associated data, are projected to the vector using linear referencing methods of the *shapely* package. To this end, each feature coordinate is assigned a value based on the normalized distance [0… 1] to its nearest coordinate point along the A/P length of the vector. The features can then be counted by dividing the vector into a user defined number of bins of equal length. The resulting data arrays enable bin-to-bin, and windowed statistical comparisons of the samples.

### Feature-to-feature distances

LAM has the option to compute pairwise Euclidean distances between nearest features (Fig S2 B). The distances can be calculated between features on one channel, e.g. DAPI, or between two channels, e.g. the distance from each Delta^+^-cell to nearest Pros^+^-cell. The features can additionally be filtered by area, volume, or another user defined variable. The algorithm finds for each feature the shortest distance to other feature in the filtered dataset.

### Feature clustering

LAM also includes an algorithm for cluster analysis that functions in a similar manner to the feature-to-feature distance calculations. Cell clusters in the midgut tend to either take the form of longer strands or a more spherical shape, and consequently defining the clusters by their shapes would be problematic. To overcome this, LAM takes the approach to cluster the cells by their proximity to each other (Fig S2 C). For each feature, LAM first finds its neighbors within a user-defined distance. Found features are then marked as a “cluster seed”. After all seeds are found, they are merged based on shared feature identification. As a result, unique clusters with no shared features are formed. The clusters can be further filtered by user-defined number of features, and are finally assigned unique cluster identification numbers.

### Gut width measurement

LAM computes the width of each midgut along its vector. The midgut is binned into segments of equal length, and nuclei with the largest distances to the vector are found. As the vector may not exactly follow the true center of the midgut, the handedness of the nuclei relative to the vector are determined. Average distance of the furthest decile of nuclei is calculated for both hand sides, and the width at each bin is the sum of these averages.

### Automatic border detection

Before running the algorithm, the nuclei area distribution is determined, and only enterocyte nuclei are included into the analysis. The borders are detected based on normalized values of (i) enterocyte nuclei distance to its nearest neighbor, (ii) midgut width, (iii) midgut width bin-to-bin difference, and (iv) enterocyte nuclei area bin-to-bin difference (default setting variables). These variables have region specific variation along the midgut’s A/P-axis, and local changes corresponds to the major region borders. In order to find local changes of the variables, a fitted fifth degree Chebyshev polynomial is subtracted from the values. To this end, for each bin (x) in the full range of bins [0… a], a total score is calculated by summing the weighted (w) deviations of each variable’s (v_i…n_) normalized value from the fitted curve (*c*):

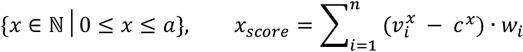

The resulting score arrays are then rescaled [0, 1] and peak detection is performed to find group average scores. To increase resolution, the border detection algorithm is run by twice the number of bins set by the user.

### Statistics

LAM includes pairwise statistical testing of control and sample groups (Fig S2 D). LAM has two types of in-built statistical testing. Firstly, bin values of sample group are tested against the respective bin of the control group resulting into a representation of p values along the A/P axis of the midgut. Secondly, total feature counts of a sample group are tested against the control group. Both tests are performed with Mann-Whitney-Wilcoxon *U* test using continuity correction. In the bin-by-bin testing, false discovery rate correction due to multiple testing is applied. Additionally, for the bin-by-bin testing, a sliding window option of user-defined size is available. The use of a sliding window has some advantages depending on input data. For example, some cell types of the midgut may be spatially too sparse for bin-to-bin testing as the cell count at each bin would be skewed towards zero. Consequently, using a sliding window to merge bins would increase the number of non-zero values in the test population, and therefore increase the strength of the statistical test.

### Drosophila stocks and husbandry

Fly stocks used in this study: w; Esg-Gal4, Tub-Gal80-ts, UAS-GFP; UAS-Flp, Act>CD2>Gal4 (Esg FO) (Jiang et al., 2009), w; Esg-Gal4, Tub-Gal80-ts, UAS-GFP (Esg^ts^) (Jiang and Edgar, 2009), Delta-LacZ (Dl-LacZ, Bloomington 11651), Gbe+Su(H)-lacZ (Su(H)-LacZ, Furriols and Bray, 2001). Flies were maintained at 25°C, on medium containing agar 0.6% (w/v), malt 6.5% (w/v), semolina 3.2% (w/v), baker’s yeast 1.8% (w/v), nipagin 2.4%, propionic acid 0.7%.

### DSS treatment

36-50 KDa DSS was obtained from Fischer Scientific (cat no. 11424352). Staged Esg FO>Delta-LacZ and Esg FO>Su(H)-LacZ pupae were collected into vials containing holidic diet (Piper et al., 2014). After eclosion the flies were kept on the holidic diet for 5 days at 18°C, and then transferred into vials containing 2% sucrose (w/v) in medium containing agar 0.5% (w/v), nipagin 2.4%, and propionic acid 0.7%, in water with or without 3% DSS, and then kept at +29°C for 5 days.

### Immunohistochemistry

For immunofluorescence staining, intestines were dissected in PBS and fixed in 8% paraformaldehyde for 3 hours. Tissues were washed with 0.1% Triton-X 100 in PBS and blocked in 1% bovine serum albumin for 1 h. Subsequently, tissues were stained with anti-β-galactosidase (1:400) (MP Biomedicals cat no: 0855976-CF) and/or anti-Prospero (1:1000) (MR1A-c, DSHB) antibodies. The samples were mounted in Vectashield mounting media with DAPI (Vector Laboratories) and imaged using the Aurox clarity confocal system (Aurox).

### Microscopy and image processing

Fixed and immunostained whole midguts were mounted in between a microscope slide with 0.12 µm spacers and a coverslip, followed by tile scan imaging by the Aurox clarity spinning disc confocal microscope from the anterior to posterior end. To reduce the image size and scanning time, stacks of only one side of the flattened midgut epithelium were obtained. For stitching the tiles and image processing in ImageJ, we generated a python script, “Stitch”, with a graphical user interface (https://github.com/hietakangas-laboratory/Stitch). “Stitch” is a programme for stitching together a series of tiff images within a directory, utilizing the ImageJ Grid/Collection plugin (Preibisch et al., 2009), and performing stitching for multiple directories in a batch process. This programme can stitch together a series of tiff images using only a companion.ome metadata file associated with the tiff series. Alternatively, as in this article, “Stitch” can utilize the tile positions output from the microscope to perform image stitching. Full usage instructions and details are available in the “Stitch” user guide. After stitching and image processing, TIFFs were converted to Imaris (Bitplane) files, and features were obtained by the Imaris spot detection algorithm (Imaris (version 9.5.1) 2019). Raw feature data, including spot surface area measurements, was exported and used as input for LAM for further analysis (see the LAM user guide for details).

## ACKNOWLEDGEMENTS

We thank Bloomington *Drosophila* Stock Center for providing fly stocks. Heini Lassila, Jorge Fuentes Rodriguez and Esa Pitkänen are acknowledged for technical assistance. We thank Paula Jouhten for critical reading of the manuscript. Funding was provided by the Academy of Finland through the MetaStem Center of Excellence funding to VH (312439) and a project grant (137530) to JM, Sigrid Juselius Foundation (VH, JM), Erkko Foundation (VH, PK), and Novo Nordisk Foundation (NNF19OC0057478 to VH). This study was facilitated by the University of Helsinki *Drosophila* core facility (Hi-Fly) and the Light microscopy unit (LMU).

## REFERENCES

Amcheslavsky, A., Jiang, J., and Ip, Y.T. (2009). Tissue Damage-Induced Intestinal Stem Cell Division in Drosophila. Cell Stem Cell 4, 49–61.

Biteau, B., Hochmuth, C.E., and Jasper, H. (2008). JNK Activity in Somatic Stem Cells Causes Loss of Tissue Homeostasis in the Aging Drosophila Gut. Cell Stem Cell 3, 442–455.

Buchon, N., Broderick, N.A., Chakrabarti, S., and Lemaitre, B. (2009). Invasive and indigenous microbiota impact intestinal stem cell activity through multiple pathways in Drosophila. Genes & Development 23, 2333–2344.

Buchon, N., Osman, D., David, F.P.A., Yu Fang, H., Boquete, J.-P., Deplancke, B., and Lemaitre, B. (2013). Morphological and Molecular Characterization of Adult Midgut Compartmentalization in Drosophila. Cell Reports 3, 1725–1738.

Dimitriadis, V.K. (1991). Fine structure of the midgut of adult Drosophila auraria and its relationship to the sites of acidophilic secretion. Journal of Insect Physiology 37, 167–177.

Dutta, D., Dobson, A.J., Houtz, P.L., Gläßer, C., Revah, J., Korzelius, J., Patel, P.H., Edgar, B.A., and Buchon, N. (2015). Regional Cell-Specific Transcriptome Mapping Reveals Regulatory Complexity in the Adult Drosophila Midgut. Cell Rep 12, 346–358.

Furriols, M., and Bray, S. (2001). A model Notch response element detects Suppressor of Hairless–dependent molecular switch. Current Biology 11, 60–64.

Gervais, L., and Bardin, A.J. (2017). Tissue homeostasis and aging: new insight from the fly intestine. Current Opinion in Cell Biology 48, 97–105.

Gillies, S., & others. (2007). Shapely: manipulation and analysis of geometric objects. Retrieved from “https://github.com/Toblerity/Shapely”

Guo, X., Yin, C., Yang, F., Zhang, Y., Huang, H., Wang, J., Deng, B., Cai, T., Rao, Y., and Xi, R. (2019). The Cellular Diversity and Transcription Factor Code of Drosophila Enteroendocrine Cells. Cell Reports 29, 4172–4185.e5.

Guo, Z., Lucchetta, E., Rafel, N., and Ohlstein, B. (2016). Maintenance of the adult Drosophila intestine: all roads lead to homeostasis. Current Opinion in Genetics & Development 40, 81–86.

Harris, C.R., Millman, K.J., van der Walt, S.J., Gommers, R., Virtanen, P., Cournapeau, D., Wieser, E., Taylor, J., Berg, S., Smith, N.J., et al. (2020). Array programming with NumPy. Nature 585, 357–362.

Hudry, B., Khadayate, S., and Miguel-Aliaga, I. (2016). The sexual identity of adult intestinal stem cells controls organ size and plasticity. Nature 530, 344–348.

Hunter, J.D. (2007). Matplotlib: A 2D Graphics Environment. Comput. Sci. Eng. 9, 90–95.

Imaris (version 9.5.1). 2019. Bitplane Inc. http://bitplane.com.

Jiang, H., and Edgar, B.A. (2009). EGFR signaling regulates the proliferation of Drosophila adult midgut progenitors. Development 136, 483–493.

Jiang, H., Patel, P.H., Kohlmaier, A., Grenley, M.O., McEwen, D.G., and Edgar, B.A. (2009). Cytokine/Jak/Stat signaling mediates regeneration and homeostasis in the Drosophila midgut. Cell 137, 1343–1355.

Jiang, H., Tian, A., and Jiang, J. (2016). Intestinal stem cell response to injury: lessons from Drosophila. Cellular and Molecular Life Sciences 73, 3337–3349.

Marianes, A., and Spradling, A.C. (2013). Physiological and stem cell compartmentalization within the Drosophila midgut. ELife 2.

McKinney, W. (2010). Data Structures for Statistical Computing in Python. (Austin, Texas), pp. 56–61.

Miguel-Aliaga, I., Jasper, H., and Lemaitre, B. (2018). Anatomy and Physiology of the Digestive Tract of *Drosophila melanogaster*. Genetics 210, 357–396.

Missiaglia, E., Jacobs, B., D’Ario, G., Di Narzo, A.F., Soneson, C., Budinska, E., Popovici, V., Vecchione, L., Gerster, S., Yan, P., et al. (2014). Distal and proximal colon cancers differ in terms of molecular, pathological, and clinical features. Annals of Oncology 25, 1995–2001.

Mowat, A.M., and Agace, W.W. (2014). Regional specialization within the intestinal immune system. Nat Rev Immunol 14, 667–685.

O’Brien, L.E. (2013). Regional Specificity in the Drosophila Midgut: Setting Boundaries with Stem Cells. Cell Stem Cell 13, 375–376.

Piper, M.D.W., Blanc, E., Leitão-Gonçalves, R., Yang, M., He, X., Linford, N.J., Hoddinott, M.P., Hopfen, C., Soultoukis, G.A., Niemeyer, C., et al. (2014). A holidic medium for Drosophila melanogaster. Nat. Methods 11, 100–105.

Preibisch, S., Saalfeld, S., and Tomancak, P. (2009). Globally optimal stitching of tiled 3D microscopic image acquisitions. Bioinformatics 25, 1463–1465.

Reiff, T., Jacobson, J., Cognigni, P., Antonello, Z., Ballesta, E., Tan, K.J., Yew, J.Y., Dominguez, M., and Miguel-Aliaga, I. (2015). Endocrine remodelling of the adult intestine sustains reproduction in Drosophila. Elife 4, e06930.

SciPy 1.0 Contributors, Virtanen, P., Gommers, R., Oliphant, T.E., Haberland, M., Reddy, T., Cournapeau, D., Burovski, E., Peterson, P., Weckesser, W., et al. (2020). SciPy 1.0: fundamental algorithms for scientific computing in Python. Nat Methods 17, 261–272.

Seabold, S., and Perktold, J. (2010). Statsmodels: Econometric and Statistical Modeling with Python. (Austin, Texas), pp. 92–96.

Strand, M., and Micchelli, C.A. (2011). Quiescent gastric stem cells maintain the adult Drosophila stomach. Proceedings of the National Academy of Sciences 108, 17696–17701.

Tian, A., Benchabane, H., Wang, Z., and Ahmed, Y. (2016). Regulation of Stem Cell Proliferation and Cell Fate Specification by Wingless/Wnt Signaling Gradients Enriched at Adult Intestinal Compartment Boundaries. PLoS Genet 12, e1005822.

van der Walt, S., Schönberger, J.L., Nunez-Iglesias, J., Boulogne, F., Warner, J.D., Yager, N., Gouillart, E., and Yu, T. (2014). scikit-image: image processing in Python. PeerJ 2, e453.

Waskom, M., Gelbart, M., Botvinnik, O., Ostblom, J., Hobson, P., Lukauskas, S., Gemperline, D.C., Augspurger, T., Halchenko, Y., Warmenhoven, J., et al. (2020). mwaskom/seaborn: v0.11.1 (December 2020) (Zenodo).

