## Supplementary figures for "LAM – an image analysis method for regionally defined organ-wide cellular phenotyping of the *Drosophila* midgut"

### Supplementary Figure 1 (related to figure 2)

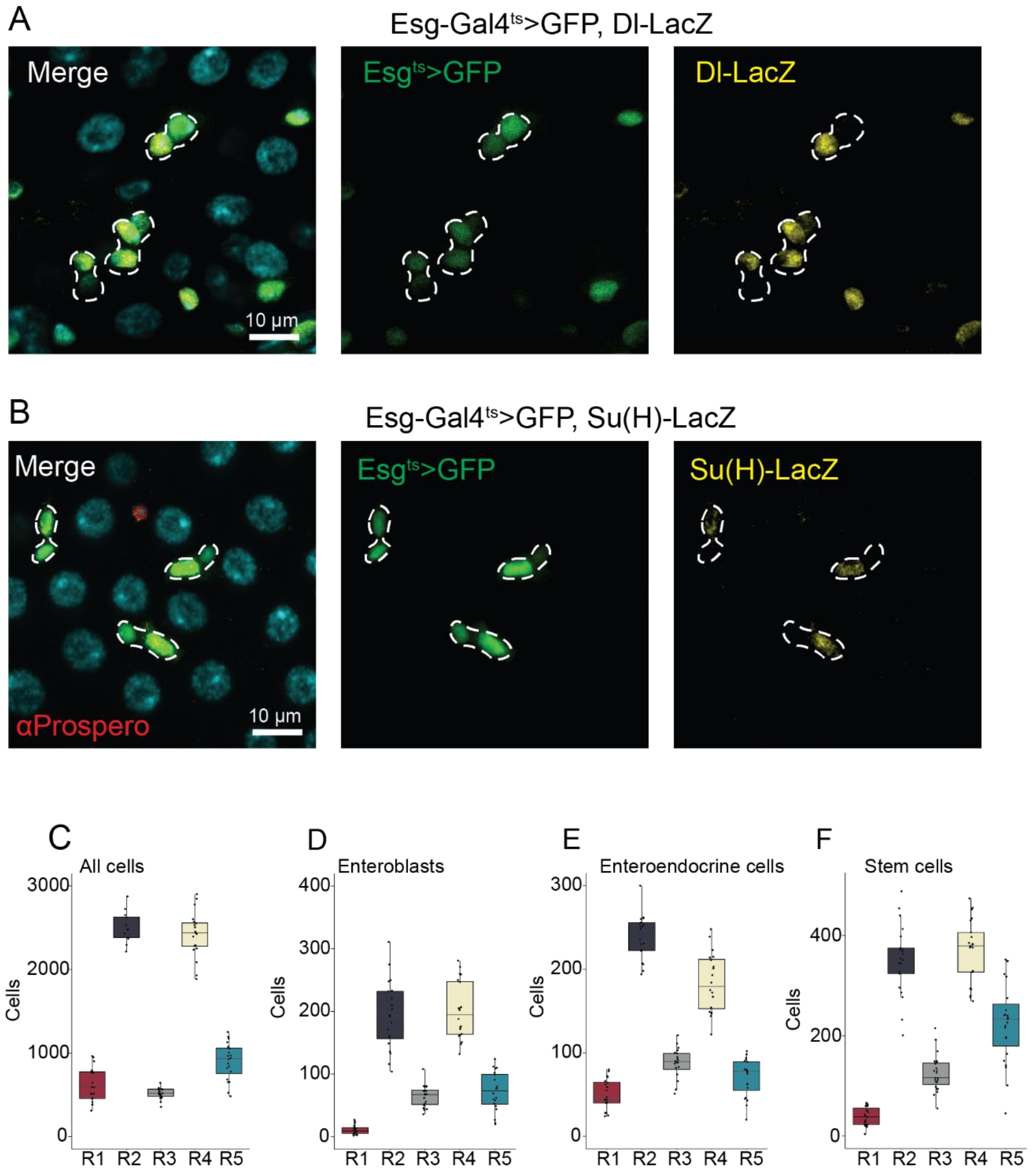

Supplementary Figure 2 (related to figures 1, 2, 3 & 4)

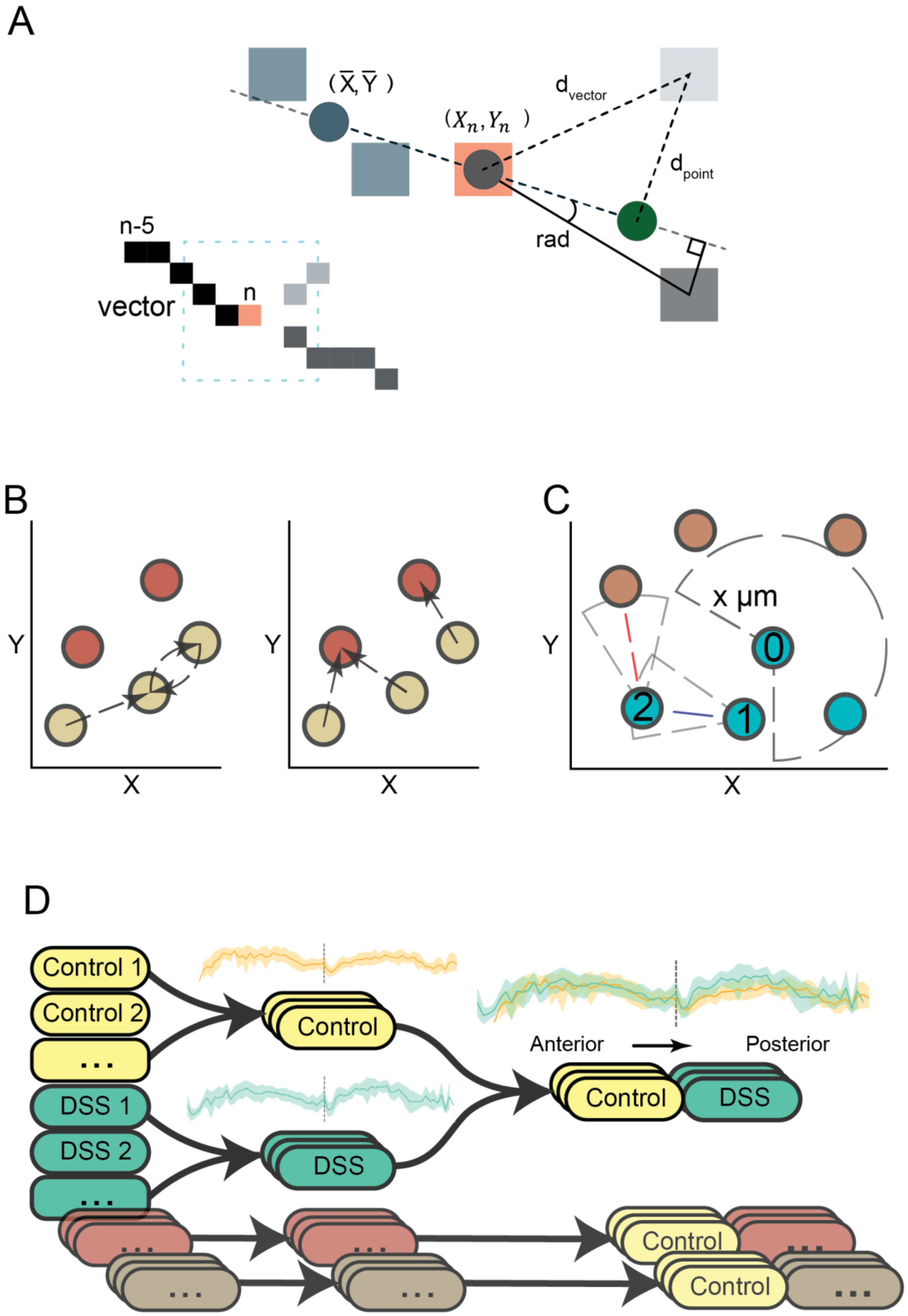

#### SUPPLEMENTARY FIGURE LEGENDS

**Supplementary Figure S1. Steady state midgut profiling (related to figure 2).**

**A & B)** Representative images of Esg-Gal4$^{ts}$ >GFP, Dl-LacZ (A) and Esg-Gal4$^{ts}$ >GFP, Su(H)-LacZ (B) stained for anti-betagalactosidase and anti-Prospero antibodies (in B only). DNA is stained with DAPI and is shown in cyan. EBs and ISCs were identified based on threshold intensity segmentation through the anti-betagalactosidase channel followed by intensity filtering through the GFP channel. EEs were identified based on threshold intensity segmentation through the anti-Prospero channel. **C-F)** Regional count of all cells (C), EBs (D) EEs (E) and ISCs (F).

**Supplementary Figure S2. Functionalities of LAM (related to figures 1, 2, 3 & 4)**

**A)** Pixel selection in skeleton vector creation. The vector is a piecewise line starting from leftmost pixels of the binary image skeleton. The vector is extended with pixel coordinates based on a scoring system that gives penalties depending on pixel's directional change and distance. With n as the last pixel of the vector, a direction giving line is formed based on coordinates of n and the average coordinate of n-1 and n-2. On this line, a projection point (green circle) is created equidistant from n as the average coordinate. For each candidate pixel, distances to n ($d_{vector}$) and the projection point ($d_{point}$) are determined, both contributing equally to the penalty. Additionally, the absolute radian changes of each pixel relative to n and the direction line is multiplied by ten, and added to the distance scores to give the full penalty. The pixel with the smallest penalty is added to the vector, and subsequently the algorithm would follow the pixels in darker grey. **B & C)** Feature-to-feature distances and clustering. Both functionalities are based on calculating distances to neighbors of each feature. **B)** Feature-to-feature distance calculations determine the nearest neighbors of a channel's features either on one channel's data set (left) or compared to a target channel's data set (right). In practice, the functionality can be used in determining cell densities and differences in cell dynamics. In the schematics, the colored circles indicate feature locations of different channels and the arrows show the nearest features in the channel that is under analysis. **C)** The clustering algorithm is based on finding neighbors of each feature on one channel to form "cluster seeds". The seeds are then merged based on shared feature ID's to form the final clusters (blue circles). In the figure, the centroid of feature number 1 falls within the cluster seed of feature 0, while feature 2 does not. However, as feature 2 is within the proximity of feature 1, during the merging of seeds all these numbered features are joined into one cluster. **D)** Pairwise sample group comparisons in LAM. All groups are first analyzed alone, and then compared against the control group. LAM analysis can include any number of sample groups, but each group is statistically tested only against the control group.
